# Resilience to persistent pain is characterized by stable periodic and increased aperiodic cortical activity

**DOI:** 10.64898/2026.04.28.721303

**Authors:** Anna M. Zamorano, Chuwen Chen, Samantha K. Millard, Boris Kleber, Peter Vuust, Herta Flor, Thomas Graven-Nielsen

## Abstract

Individual variability in pain perception raises fundamental questions about how biological and experiential factors shape pain processing. Cognitive-demanding motor training is a key driver of use-dependent brain plasticity and may contribute to differences in pain responses. Using musicians as a model of cognitive-motor expertise, we examined how such experience influences cortical dynamics and pain perception during experimentally induced prolonged musculoskeletal pain. Resting-state electroencephalography (EEG) was recorded in musicians and non-musicians before (Day 1) and during pain development (Days 3 and 8) following intramuscular nerve growth factor (NGF) administration. We parameterized periodic (alpha peak frequency, power, frontal asymmetry) and aperiodic (exponent, offset) components of the EEG signal to characterize intrinsic cortical activity. During pain development, non-musicians exhibited slowing of peak alpha frequency, a neural marker associated with ongoing pain. In contrast, musicians showed preserved alpha dynamics and greater left frontal asymmetry, reflecting resilient top-down pain regulation. Musicians also displayed higher aperiodic exponent across sessions, suggesting that musical training shapes the excitation-to-inhibition (E:I) balance potentially reflecting a shift toward greater inhibitory activity. Notably, across all participants, only aperiodic features improved the prediction of pain severity, with higher exponents and higher offsets associated with lower pain ratings. These findings demonstrate that cognitive-motor training shapes cortical dynamics during sustained pain, supporting more stable, resilient cortical responses to pain. Such training also contributes to inter-individual variability in pain processing. Moreover, this study identifies aperiodic EEG components as predictors of pain severity and resilience.

## BACKGROUND

Pain varies considerably between individuals, as reflected in both behavioral and neural responses to identical nociceptive input [5]. This variability is shaped by biological, cognitive, and psychosocial factors [5,13]. Increasing evidence suggests that experiential factors, particularly cognitively demanding motor activities such as musical training, may influence pain responses [30,53,55,56]. Musical practice induces use-dependent plasticity in the brain activity, including enhanced cortical excitability [26,41], connectivity [32,51], and inhibitory control [9,24,28], processes that are also implicated in pain modulation. Musicians exhibit heightened sensitivity and increased cortical responses to phasic pain [53,57], yet report lower pain intensity and interferences and have preserved corticospinal excitability (CE) during long-term pain [55,56]. These adaptations scale with training duration [53,55,56], positioning musicians as a valuable model for studying how long-term cognitive-motor experience and its associated use-dependent plasticity shapes individual pain responses.

Chronic pain has been linked to thalamocortical dysrhythmia, an abnormal slowing of thalamocortical oscillations [47]. In particular, slower peak alpha frequency (PAF) in resting-state electroencephalographic (rs-EEG) recordings has been associated with experimental [4] and acute postoperative pain intensity [23,35], as well as chronic pain presence [29,47,49]. Additionally, slowing of PAF has been observed during experimental pain [6,15,34]. Changes in alpha power (AP) have also been reported in the context of pain, with some studies showing decreases, interpreted as reduced cortical inhibition [47], and others reporting increases [10,11,29], potentially reflecting compensatory inhibitory processes or abnormalities of neural resources allocation in the resting brain [10]. Right frontal alpha asymmetry (FAA), an index of affective states, has been associated with maladaptive responses such as helplessness and pain avoidance [43]. Together, these alpha-band features may represent neurophysiological markers of individual differences in pain responsiveness and neuroadaptability to pain.

Beyond oscillatory activity, EEG also captures non-oscillatory, aperiodic dynamics (exponent and offset) distributed across all frequency bands [8]. Parameterization of these components provides additional information about underlying neural population activity: the spectral exponent reflects cortical excitation-inhibition balance [20], while offset represents population-level neural firing rates [33]. Despite these features having gained increasing attention as markers of cortical state, aperiodic components remain largely unexplored in the context of motor training, pain processing, and their potential interaction.

To address this gap, we investigated how prior, long-term cognitive-motor training influences resting-state EEG markers associated with sustained musculoskeletal pain. Musicians and non-musicians underwent experimental induction of prolonged muscle pain using intramuscular nerve growth factor (NGF), while rs-EEG was recorded before (Day 1) and during pain development (Day 3 and 8). We parameterized both oscillatory (AP, PAF, FAA) and aperiodic (exponent, offset) components of the EEG spectrum to characterize intrinsic cortical dynamics. We hypothesized that prior motor experience in musicians would shape EEG markers at baseline (Day 1), manifested as faster PAF, decreased AP, and left-sided FAA, compared to non-musicians. As long-term motor training induces lasting experience-dependent corticomotor adaptations [55], we further expected these features to remain stable in musicians during pain (Day 3, Day 8), while non-musicians would show pain-related changes (slowing PAF, AP reduction, right FAA). We further conducted a posthoc analysis of aperiodic EEG components (exponent, offset) to broaden spectral characterization and tested whether any EEG feature predicted pain severity.

## METHODS

### Participants

Thirty-nine participants were recruited via adverts at Aalborg University, Aarhus University, and The Royal Academy of Music, Aarhus/Aalborg. Nineteen of these were healthy musicians (6 females, mean age 25.0 ± 2.6 years) consisting of 9 amateurs (4,406 ± 2,776 hours of accumulated training and 8.1 ± 3.7 hours of weekly training) and 10 conservatory-trained musicians (15,540 ± 6,621 hours of accumulated training and 27.6 ± 14.2 hours of weekly training). The control group included 20 healthy non-musicians (nine female, 19 right-handed, mean age 26.9 ± 5.3 years) without any prior formal or informal music training.

This study is part of a larger longitudinal project investigating the interaction between experimental muscle pain and prior use-dependent plasticity associated with sensorimotor training (ClinicalTrials.gov–NCT04457466). As part of this comprehensive project, a broad range of measures were collected and previously reported [53,54], including psychological variables, neurophysiological responses, experimentally-induced pain ratings and pain distribution, quantitative sensory assessments, as well as corticomotor excitability measured using transcranial magnetic stimulation (TMS). Additional data such as resting-state electroencephalography (EEG) and somatosensory evoked potentials were also collected. In the context of this large dataset, the present manuscript focuses exclusively on the outcomes related to resting-state EEG, which have not been previously published. In addition, this manuscript examines the relationship between the resting-state EEG periodic and aperiodic features and the prolonged muscle pain experience, with specific emphasis on how these dynamics relate to musical experience. The analyses focusing on unpublished resting-state EEG outcomes and their relation to pain were pre-registered (osf.io/4c2b8). The study was approved by the local ethics committee (N-20170040) and aligned with the Declaration of Helsinki.

Exclusion criteria were neurological, cardiorespiratory, or mental disorders, or pregnancy, as well as history of chronic pain or current acute pain. The sample size was estimated using G*Power [12] and based on previous publications using a similar approach [6]. This was done to ensure 80% power for detecting at least a medium-sized interaction effect (*η*^*2*^ ≥ 0.06) between *group* and *time* on peak alpha frequency, using a mixed-model analysis of variance (ANOVA) at an alpha level of 0.05. The sample size required for each primary outcome (nociceptive evoked potentials and corticomotor maps, published elsewhere [53,54], and the EEG features of the current publication) were calculated separately, and the larger of all sample sizes was chosen to ensure sufficient power for all outcomes. All participants received written and verbal information about the study and provided written consent.

### Experimental Procedure

The experiment involved 3 sessions (Day 1, Day 3, and Day 8) over 8 days. Participants were seated in a comfortable chair for the laboratory sessions. At the end of Day 1, all participants received an injection of NGF into the right first dorsal interosseous (FDI) muscle to induce prolonged muscle pain in the hand for several days. At the beginning of each session, participants reported the demographic (Day 1) and the pain-related questionnaires (Day 1, Day 3, Day 8). Subsequently, on Day 1 (before the NGF injection), Day 3, and Day 8, cortical activity at rest and with eyes open was registered by using EEG.

### Experimental Prolonged Muscle Pain

Muscle pain and hyperalgesia were induced by intramuscular injections of NGF into the right FDI muscle after all assessments on Day1. Sterile solutions of recombinant human beta-NGF were prepared by the pharmacy (Skanderborg Apotek, Denmark). The site of injection was cleaned with alcohol, and NGF solution (5μg/0.5 mL) was immediately injected into the FDI of the right hand.

Pain ratings in resting conditions were reported at the beginning of Day 3 and Day 8 using a Numeric Rating Scale (NRS) with 0 defined as “no pain” to 10 corresponding to “worst pain imaginable”. In addition, NRS ratings for the worst, the least, and the average NGF-induced muscle pain during the entire week were reported on Day 8. The NRS pain ratings reported in this study are described in detail in Zamorano et al. [53].

### Electrophysiological activity

Electroencephalographic (EEG) activity was recorded using an active electrode cap (g.SCARABEO, g.tec, Medical Engineering GmbH, Austria). The electrode montage included 64 electrodes consisting of the modified 10-20 system with a left earlobe (A1) reference. The ground electrode was placed at position AFz. The impedance of all electrodes was kept below 20 kΩ and assessed by the EEG system software (g.Recorder, g.tec, Medical Engineering GmbH, Austria). During recordings, the EEG signals were amplified and digitized using a sampling rate of 1200 Hz (g.Hlamp, g.tec, Medical Engineering GmbH, Austria). During the entire EEG recording, participants were seated in a comfortable chair in a quiet room and instructed to remain awake and relaxed. The resting-state EEG recordings lasted approximately 4 minutes (262.793 ± 44.61 seconds).

EEG data preprocessing and analysis were conducted in MATLAB 2024b (The Mathworks, Inc) using the openly available DISCOVER-EEG pipeline version 2.0.0 [21], with modifications to specific preprocessing steps, alongside the EEGLAB (v2022.0) and FieldTrip (20231220) toolboxes [7,36] for additional outcome. The automatic preprocessing steps included down sampling to 250 Hz, rejection of bad channels, high pass filter with a default transition band of 0.25Hz to 0.75Hz, re-referencing to the average reference, notch filtering at 49–51 Hz (note that this replaced *pop_cleanline()* from the automatic pipeline), independent component analysis (ICA), removal of artifactual components, interpolation of rejected channels, and exclusion of bad time segments, and segmentation into 2-second epochs with a 50% overlap. Power spectra were computed between 1-100Hz using the Slepian multitaper method with ± 1Hz frequency smoothing. Data from each EEG channel was used to produce a single power spectrum estimate per channel. For all analyses, the alpha frequency band was defined as 8–12.9 Hz. Global average alpha power was computed by applying a 10log10 transformation to each channel, followed by averaging across all channels. Peak alpha frequency (PAF) was computed based on the global power spectrum within the alpha range by calculating the center of gravity of the power spectral density within the DISCOVER-EEG pipeline. Regarding additional measures not using the DISCOVER-EEG pipeline, frontal alpha asymmetry (FAA) was calculated as the difference in alpha power between the right (F4) and left (F3) frontal electrodes (FAA = Alpha Power at F4 − Alpha Power at F3) [43]. We also used FieldTrip’s ft_freqanalysis function over the 1–40 Hz frequency range to isolate aperiodic from oscillatory activity. The aperiodic component of the EEG signal (1/f activity) was characterized using a conventional parametrization approach called FOOOF (Fitting Oscillations & One-Over-F)[8], extracting the exponent and offset from the power spectrum. The exponent is expressed as the slope of the spectrum in log–log space, yielding negative values, with less negative (flatter) slopes indicating relatively greater high-frequency power, whereas the offset reflects the broadband power (i.e., a vertical shift of the spectrum across frequencies).

### Statistical Analysis

Data are presented as means and standard deviations in Table 1 and Figure 1. All analyses were conducted using IBM SPSS Statistics 29 for Windows. Prior to analysis, data were screened for assumptions of normality, sphericity, homogeneity of variance, and independence of errors using descriptive plots and standard statistical tests.

**Table 1.** Descriptive statistics of EEG features across groups and sessions. Values are presented as mean ± standard deviation.

| EEG-feature | Group | Day 1 | Day 3 | Day 8 |
| --- | --- | --- | --- | --- |
| Peak Alpha Frequency | Musicians | 10.37 ± 0.33 | 10.34 ± 0.35 | 10.32 ± 0.40 |
| | Non-musicians | 10.28 ± 0.33 | 10.12 ± 0.38‡ | 10.20 ± 0.37 |
| Alpha Power (dB) | Musicians | -10.49 ± 2.63 | -9.57 ± 3.15 | -9.88 ± 3.98§ |
| | Non-musicians | -10.13 ± 3.68 | -9.39 ± 4.16 | -9.34 ± 3.98§ |
| Frontal Alpha Asymmetry | Musicians | 0.10 ± 0.65 | 0.05 ± 0.78 | 0.58 ± 0.51†$\Phi$ |
| | Non-musicians | 0.23 ± 0.86 | -0.07 ± 0.78 | -0.01 ± 0.87Φ |
| Offset | Musicians | -0.25 ± 0.18 | -0.24 ± 0.17 | -0.24 ± 0.17 |
| | Non-musicians | -0.36 ± 0.27 | -0.32 ± 0.28 | -0.31 ± 0.29 |
| Exponent | Musicians* | 1.27 ± 0.15 | 1.27 ± 0.12 | 1.27 ± 0.13 |
| | Non-musicians* | 1.13 ± 0.22 | 1.18 ± 0.20 | 1.17 ± 0.20 |

‡Significantly lower than Day 1 within non-musicians ( $p < .001$ )
§ Significantly higher than Day 1 across all participants (main effect of Time, $p < .05$ ).
† Significantly higher than Day 1 and Day 3 within musicians ( $p < .05$ ).
$\Phi$ Significant difference between groups within the same day ( $p < .05$ ).
\* Significantly higher in musicians than non-musicians across sessions (main effect of Group, $p < .05$ ).

**Figure 1.**
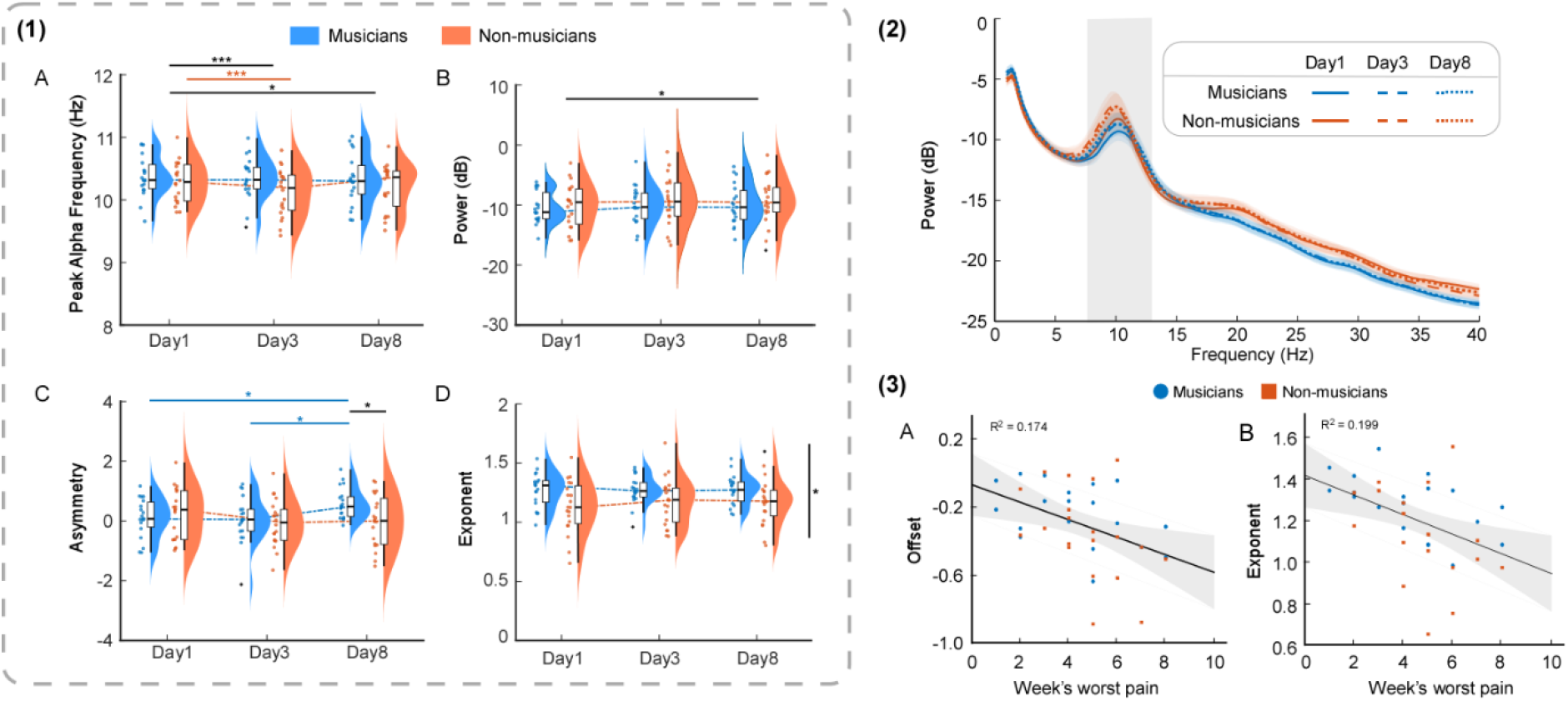
Electrophysiological markers in musicians (blue) and non-musicians (orange) and their relationship with pain. **1:** Distribution and group comparisons of EEG features across sessions (Day 1, 3, and 8): (A) Peak alpha frequency showed a reduction over time in non-musicians but relative stability in musicians. (B) Alpha power increased during pain development, with a significant rise from Day 1 to Day 8 across participants. (C) Frontal alpha asymmetry showed a greater leftward asymmetry in musicians compared to non-musicians, particularly at Day 8. (D) Aperiodic exponent was consistently higher in musicians across sessions, indicating a change in the excitation-to-inhibition (E:I) balance. **2:** Grand-average power spectra across sessions and groups. **3:** Associations between baseline (Day 1) aperiodic parameters and pain severity (week’s worst pain). (A) Aperiodic offset and (B) exponent both show negative relationships with pain severity, indicating that flatter spectra and higher offset are associated with lower perceived pain.

Periodic (PAF, AP, FAA) and aperiodic (exponent, offset) EEG features were compared across Time (Day1, Day3, Day8) and Group (musicians vs. non-musicians) using two-way repeated-measures ANOVAs. Significant main effects or interactions were followed up with Bonferroni-corrected post hoc comparisons.

Pearson correlations were computed to examine whether EEG features (PAF, AP, FAA, exponent, offset) measured during pain-free conditions (Day 1) were associated with the peak of NGF-induced pain (Day3) and with week’s worst pain, both across all participants and within groups.

To determine whether baseline (Day 1) EEG features predicted subsequent pain outcomes, two hierarchical multiple regressions were conducted. Each regression predicted (1) Day3 pain intensity or (2) week’s worst pain. In Step 1, periodic EEG components (PAF, AP) were entered. In Step 2, aperiodic components (exponent, offset) were added to evaluate their incremental predictive value. Overall model significance was assessed using F-tests, and improvements in explained variance were evaluated using F-change statistics.

For all tests used, the level for statistically significant difference was set at *p* < 0.05. Bonferroni’s procedure was used to correct for multiple comparisons.

## RESULTS

### Slowing of peak alpha frequency in non-musicians but not in musicians

A significant *Time × Group* interaction was observed (*F*(2, 74) = 3.258, *p* = .044, *η*^*2*^*p* = 0.08), with post hoc comparisons indicating a stronger decline in non-musicians, particularly on Day3 compared with Day1 (*p* < .001, *d* = 0.44). No significant changes across sessions were observed in musicians (all *p* >.468, all *d* < 0.15). Main effects of *Time* indicated that Peak alpha frequency (Table 1, Fig. 1A) decreased across sessions (*F*(2, 74) = 6.446, *p* = .003, *η*^*2*^*p* = 0.15). Post hoc comparisons showed lower values on Day3 (*p* = .001, *d* = 0.26) and Day8 (*p* = .047, *d* = 0.19) compared with Day1. The main effect of *Group* was not statistically significant, *F*(1, 37) = 1.685, *p* = .202, *η*^*2*^*p* = 0.04.

### Alpha power increases over time

A main effect of *Time* was observed, where alpha power (Table 1, Fig. 1B) increased across sessions (*F*(2, 74) = 4.084, *p* = .021, *η*^*2*^*p* = 0.10). Post hoc comparisons showed higher values on Day8 compared with Day1 (*p* = .05, *d* = 0.20). No significant main effect of *Group (F*(1, 37) = 0.199, *p* = .658, *η*^*2*^*p* < 0.01), or *Time × Group* interaction was observed (*F*(2, 74) = 0.89, *p* = .915, *η*^*2*^*p* < 0.01).

### Increases in left frontal alpha asymmetry in musicians

Frontal alpha asymmetry (Table 1, Fig. 1C) showed a significant Time × Group interaction (*F*(2, 74) = 3.723, *p* = .029, *η*^*2*^*p* = 0.09). Musicians exhibited greater left-sided FAA on Day8 compared with non-musicians (*p* = .014, *d* = 0.79). Within the musician group, FAA increased on Day8 compared with both Day1 (*p* = .049, *d* = 0.63) and Day3 (*p* = .011, *d* = 0.71). No significant changes were observed in non-musicians (all *p* >. 484, *d* < 0.40). No significant main effects of *Group* (*F*(1, 37) = 0.109, *p* = .299, *η*^*2*^*p* = 0.03), or *Time* (*F*(2, 74) = 1.109, *p* = .299, *η*^*2*^*p* = 0.03) were found.

### Higher aperiodic exponent in musicians

Aperiodic exponent (Table 1, Fig. 1D) was significantly higher in musicians than in non-musicians (main effect of *Group: F*(1, 37) = 4.527, *p* = .040, *η*^*2*^*p* = 0.11). No significant effects of *Time* (*F*(2, 74) = 1.022, *p* = .365, *η*^*2*^*p* = 0.03) or *Time × Group* interaction (*F*(2, 74) = 1.079, *p* = .345, *η*^*2*^*p* = 0.03) were observed.

For the aperiodic offset, no significant interaction *Time × Group* (*F*(2, 74) = 0.563, *p* = .572, *η*^*2*^*p* = 0.01) or main effects of *Group* (*F*(1, 37) = 1.511, *p* = .227, *η*^*2*^*p* = 0.04) or *Time* were observed (*F*(2, 74) = 1.074, *p* = .227, *η*^*2*^*p* = 0.04). This suggests stability of the aperiodic offset across sessions and groups.

### Pain intensity correlates with aperiodic but not periodic EEG features

Contrary to our hypothesis, baseline periodic EEG measures (PAF, AP, and FAA) were not significantly correlated with pain intensity on any day (all p>.05).

In contrast, baseline aperiodic parameters showed significant associations with pain ratings across participants (Fig. 3A–B). Higher offsets were associated with lower worst pain ratings during the week (*r* = –.417, *p* = .008). Higher exponents (i.e., a steeper spectral slope) values were associated with lower pain ratings on Day3 (*r* = –.403, p = .011), Day8 (*r* = –.338, *p* = .035), and lower worst pain ratings during the week (*r* = –.446, *p* = .004).

Subgroup analyses indicated that these associations were driven by the musician group, where exponent correlated negatively with Day3 pain ratings (*r* = –.494, *p* = .032) and worst pain during the week (*r* = –.554, *p* = .014). In non-musicians, no significant correlations were observed; however, a similar negative trend between exponent and worst pain ratings was evident (*r* = –.379, *p* = .099).

The offset was not significantly associated with pain outcomes in either group, although comparable negative associations with worst pain were observed in both musicians (*r* = –.424, *p* = .070) and non-musicians (*r* = –.423, *p* = .063).

### Aperiodic EEG features predict pain severity

Two hierarchical regression analyses examined whether baseline EEG measures predicted pain intensity on Day3 and worst pain during the week. In the first step, periodic EEG measures (PAF, AP) were entered as predictors. This model was not significant for pain ratings on Day 3 (*F*(2,36) = 0.88, *p* =.423, *R*^*2*^*=*.047) or for the week’s worst pain (*F*(2,36) = 0.17, *p* = .843, *R*^2^=.009).

In the second step, aperiodic components (exponent, offset) were added to the model. This significantly improved model fit for Day 3 pain ratings (*ΔF*(2,34) = 4.30, *p* = .022, ΔR^2^ = 0.193) and for worst weekly pain (*ΔF*(2,34) = 4.50, *p* = .019, *ΔR*^*2*^ = .207). The final model explained 24% of the variance in Day 3 pain (*F*(4,34) = 2.67, *p* = .048, *R*^*2*^ *=* 0.239), and 21.7% of the variance of the worst pain during the week. However, the model for worst pain did not reach statistical significance (*F*(4,34) = 2.35, *p* = .074, *R*^*2*^ = .217). No individual predictor was significant in either model (all *p* > 0.05).

## DISCUSSION

In the present study, we found that prolonged musculoskeletal pain led to a pronounced slowing of peak alpha frequency in non-musicians, highlighting the effects of persistent pain on alpha rhythms. Meanwhile, musicians exhibited more stable oscillatory dynamics and stronger left-sided frontal alpha asymmetry during sustained pain, suggesting more stable and resilient cortical responses to pain. Musicians also showed higher aperiodic exponent values compared to non-musicians and independent of pain, suggesting that cognitive-motor training may shape the excitation-inhibition (E:I) balance, potentially reflecting a shift toward greater inhibitory activity. Contrary to our hypothesis, baseline PAF and alpha power did not predict pain severity. However, the addition of aperiodic EEG parameters (exponent and offset) contributed to the prediction of pain severity, suggesting that, when an individual has a more excitable cortical state at baseline—indexed by higher offset and steeper slope—lower pain intensity is experienced.

### Evidence for neural pain resilience in musicians

The present work is partly consistent with previous work linking slower PAF and slowing of PAF to experimental pain [4,6,16,46], postoperative [23,35] and chronic pain states [3,11,47,49]. However, PAF slowing was only observed in non-musicians, who showed a marked reduction in PAF shortly after pain induction. Slowing of PAF has been interpreted as a marker of altered thalamocortical dynamics and reduced efficiency of top-down regulatory processes during sustained nociceptive input [4,10,31]. Thus, the relative stability of PAF in musicians may therefore reflect greater resilience of intrinsic cortical dynamics during pain or efficiently regulated top-down control, potentially shaped by goal-directed cognitive-motor training [1,26,56].

This interpretation is supported by converging evidence from pain ratings and corticomotor excitability (CE) measures collected as part of the same project [53,55]. In non-musicians, NGF induced a reduction in CE across days and higher pain ratings, whereas musicians showed stable CE profiles and lower pain scores. Moreover, in musicians, corticospinal parameters were associated with both weekly and accumulated training, reinforcing the notion that motor experience counteracts pain-related plasticity. Collectively, prior cognitive-motor experience may shape pain responses and contribute to pain resilience, likely through enduring forms of experience-dependent plasticity that stabilize neural function and behavior [1,22,56] .

Alpha power increased over time, with higher power on Day 8 compared to Day 1. This progressive increase in alpha power may reflect enhanced inhibitory gating and reduced cortical excitability during sustained nociceptive input, suggesting that cortical networks gradually engage regulatory mechanisms to manage ongoing pain [10]. This effect was not specific to musicians, suggesting it reflects a general neural pain-related adaptation rather than a motor experience-dependent modulation.

However, the groups differed in frontal alpha asymmetry (FAA) during sustained pain. Musicians showed greater left-frontal alpha asymmetry over time (i.e., positive values), a pattern associated with approach-related motivational states and adaptive coping responses [43]. FAA reflects the relative difference in alpha power between left and right frontal regions and provides an indirect index of hemispheric differences in cortical activity, with positive values indicating relatively greater left frontal cortical activity [44]. Greater relative left-FAA is associated with approach-related behavior and lower pain-related catastrophizing [27,43,59]. In this context, the leftward shift in FAA observed in musicians may reflect sustained engagement in goal-directed behavior despite discomfort [45,56]. This is consistent with the demands of intensive musical training, which may foster approach motivation and buffer against affective withdrawal in response to pain [53,55,56].

### Increased aperiodic exponent in musicians

In the present study, musicians also exhibited higher aperiodic exponent values compared with non-musicians. Moreover, exponent values were not modulated by pain across sessions, suggesting that these differences likely reflect stable group characteristics rather than transient pain-related neural changes. The aperiodic exponent provides an index of the slope of the neural power spectrum and is thought to reflect large-scale properties of cortical population activity [8,33,48]. Variations in the exponent have been proposed to relate to shifts in excitation–inhibition (E:I) balance, with steeper slopes interpreted as reflecting relatively greater inhibitory contributions [20]. Aperiodic activity has been shown to vary with cognitive demands, including working memory and response inhibition [8,50,58]. Notably, steeper spectral slopes have been observed during controlled inhibition, suggesting a link between aperiodic dynamics and cognitive control mechanisms [38,39,58]. In this context, higher exponent values in musicians may relate to neural processes supporting inhibitory control shaped by long-term cognitive-motor training.

Converging evidence from neurophysiological studies further supports this interpretation, as musicians have been reported to exhibit reduced corticomotor excitability, potentially reflecting long-term adaptations in inhibitory mechanisms associated with extensive motor training [37,52,54]. Given that musical training involves extensive and sustained practice requiring precise and coordinated actions, group differences in exponent may reflect long-term adaptations of cortical networks supporting motor-skill acquisition and cognitive expertise [18,19,25,26,41].

### Aperiodic components predict pain intensity

Interestingly, contrary to our hypothesis and previous literature reporting associations between pain-free alpha features, particularly alpha power and peak alpha frequency, with pain severity [2,4,14,15,17,35,40], periodic features in the present study did not predict subsequent pain severity. Instead, aperiodic parameters measured during the pain-free baseline were associated with ongoing pain ratings across participants and significantly improved the prediction of pain severity. Specifically, higher pain-free baseline exponent and higher offset values were associated with lower pain severity.

Because the exponent and offset capture broadband properties of neural population activity and excitation–inhibition balance [8,20,48], they may index baseline cortical gain states that shape how sensory information is processed. A steeper spectral slope (i.e., higher exponent values) has been linked to a relative increase in lower-frequency activity compared to higher frequencies and, consequently, to shifts in the balance between excitatory and inhibitory neural activity [20,42,48]. Individuals with steeper power spectra and higher baseline offset values may therefore exhibit neural dynamics that limit the amplification of nociceptive signals, resulting in lower perceived pain. Importantly, these findings highlight the potential relevance of aperiodic EEG parameters as neurophysiological markers of pain susceptibility and extend previous work by demonstrating that non-oscillatory components of neural activity may provide predictive information about pain severity beyond traditional periodic EEG markers [23,40].

### Limitations

Although the observed differences are consistent with long-term sensorimotor adaptations associated with musical expertise, the cross-sectional comparison between musicians and non-musicians does not allow causal inference regarding the effects of musical training. Longitudinal designs examining neural changes across the course of musical training would be necessary to establish causality. Additionally, we did not account for other factors, such as lifestyle factors, coping strategies, or attention, which could have shed light on whether the observed differences in periodic and aperiodic EEG activity are linked to top-down control of pain behaviour. Although the present design focused on pain-free baseline EEG as a predictor of pain severity, future studies with larger samples and additional behavioral or physiological measures will be important to better account for these sources of variability and refine the potential use of EEG markers as predictors of pain vulnerability.

## CONCLUSION

This study shows that repetitive cognitive-motor training, as exemplified by musical practice, supports more stable, resilient cortical responses to pain. Musicians also exhibited higher aperiodic exponent values, indicating differences in intrinsic cortical dynamics that may reflect training-related adaptations. These effects are likely due to strengthening of synaptic connections, where experience-dependent plasticity associated with cognitive-motor training stabilizes neural function and behavior [22]. This provides a mechanistic basis for how cognitive-motor training may shape brain responses to pain. Notably, aperiodic EEG features recorded before pain induction improved the prediction of pain severity, underscoring the importance of incorporating these measures into investigations of pain processing. Future work should investigate whether such training-induced effects offer resilience against the maladaptive effects of chronic pain or could aid recovery in clinical settings.

## References

[1] Altenmüller E, Furuya S. Brain Plasticity and the Concept of Metaplasticity in Skilled Musicians. In: Laczko J, Latash ML, editors. Progress in Motor Control: Theories and Translations. Advances in Experimental Medicine and Biology. Cham: Springer International Publishing, 2016. pp. 197–208. doi:10.1007/978-3-319-47313-0_11.

[2] Babiloni C, Brancucci A, Percio CD, Capotosto P, Arendt-Nielsen L, Chen ACN, Rossini PM. Anticipatory Electroencephalography Alpha Rhythm Predicts Subjective Perception of Pain Intensity. The Journal of Pain 2006;7:709–717.

[3] Cavaleri R, McLain NJ, Heindel M, Schrepf A, Rodriguez LV, Kutch JJ. Peak alpha frequency is related to the degree of widespread pain, but not pain intensity or duration, among people with urologic chronic pelvic pain syndrome. PAIN Reports 2025;10:e1251.

[4] Chowdhury NS, Bi C, Furman AJ, Chiang AKI, Skippen P, Si E, Millard SK, Margerison SM, Spies D, Keaser ML, Da Silva JT, Chen S, Schabrun SM, Seminowicz DA. Predicting Individual Pain Sensitivity Using a Novel Cortical Biomarker Signature. JAMA Neurology 2025;82:237–246.

[5] Coghill RC, McHaffie JG, Yen YF. Neural correlates of interindividual differences in the subjective experience of pain. Proceedings of the National Academy of Sciences of the United States of America 2003;100:8538–8542.

[6] De Martino E, Gregoret L, Zandalasini M, Graven-Nielsen T. Slowing in Peak-Alpha Frequency Recorded After Experimentally-Induced Muscle Pain is not Significantly Different Between High and Low Pain-Sensitive Subjects. The Journal of Pain 2021;22:1722–1732.

[7] Delorme A, Makeig S. EEGLAB: An open source toolbox for analysis of single-trial EEG dynamics including independent component analysis. Journal of Neuroscience Methods 2004;134:9–21.

[8] Donoghue T, Haller M, Peterson EJ, Varma P, Sebastian P, Gao R, Noto T, Lara AH, Wallis JD, Knight RT, Shestyuk A, Voytek B. Parameterizing neural power spectra into periodic and aperiodic components. Nat Neurosci 2020;23:1655–1665.

[9] Fasano MC, Semeraro C, Cassibba R, Kringelbach ML, Monacis L, de Palo V, Vuust P, Brattico E. Short-Term Orchestral Music Training Modulates Hyperactivity and Inhibitory Control in School-Age Children: A Longitudinal Behavioural Study. Front Psychol 2019;10. doi:10.3389/fpsyg.2019.00750.

[10] Fauchon C, Kim JA, El-Sayed R, Osborne NR, Rogachov A, Cheng JC, Hemington KS, Bosma RL, Dunkley BT, Oh J, Bhatia A, Inman RD, Davis KD. A Hidden Markov Model reveals magnetoencephalography spectral frequency-specific abnormalities of brain state power and phase-coupling in neuropathic pain. Commun Biol 2022;5:1000.

[11] Fauchon C, Kim JA, El-Sayed R, Osborne NR, Rogachov A, Cheng JC, Hemington KS, Bosma RL, Dunkley BT, Oh J, Bhatia A, Inman RD, Davis KD. Exploring sex differences in alpha brain activity as a potential neuromarker associated with neuropathic pain. PAIN 2022;163:1291.

[12] Faul F, Erdfelder E, Buchner A, Lang A-G. Statistical power analyses using G*Power 3.1: tests for correlation and regression analyses. Behav Res Methods 2009;41:1149– 1160.

[13] Fillingim RB. Individual differences in pain: understanding the mosaic that makes pain personal. PAIN 2017;158:S11.

[14] Furman AJ, Meeker TJ, Rietschel JC, Yoo S, Muthulingam J, Prokhorenko M, Keaser ML, Goodman RN, Mazaheri A, Seminowicz DA. Cerebral peak alpha frequency predicts individual differences in pain sensitivity. NeuroImage 2018;167:203–210.

[15] Furman AJ, Prokhorenko M, Keaser ML, Zhang J, Chen S, Mazaheri A, Seminowicz DA. Prolonged Pain Reliably Slows Peak Alpha Frequency by Reducing Fast Alpha Power. eLife 2024;13. doi:10.7554/eLife.102096.1.

[16] Furman AJ, Prokhorenko M, Keaser ML, Zhang J, Chen S, Mazaheri A, Seminowicz DA. Prolonged Pain Reliably Slows Peak Alpha Frequency by Reducing Fast Alpha Power. 2021:2021.07.22.453260. doi:10.1101/2021.07.22.453260.

[17] Furman AJ, Thapa T, Summers SJ, Cavaleri R, Fogarty JS, Steiner GZ, Schabrun SM, Seminowicz DA. Cerebral peak alpha frequency reflects average pain severity in a human model of sustained, musculoskeletal pain. Journal of Neurophysiology 2019;122:1784–1793.

[18] Furuya S, Altenmüller E. Flexibility of movement organization in piano performance. Frontiers in Human Neuroscience 2013;7:173.

[19] Furuya S, Oku T, Nishioka H, Hirano M. Surmounting the ceiling effect of motor expertise by novel sensory experience with a hand exoskeleton. Science Robotics 2025;10:eadn3802.

[20] Gao R, Peterson EJ, Voytek B. Inferring synaptic excitation/inhibition balance from field potentials. NeuroImage 2017;158:70–78.

[21] Gil Ávila C, Bott FS, Tiemann L, Hohn VD, May ES, Nickel MM, Zebhauser PT, Gross J, Ploner M. DISCOVER-EEG: an open, fully automated EEG pipeline for biomarker discovery in clinical neuroscience. Sci Data 2023;10:613.

[22] Griffith EC, West AE, Greenberg ME. Neuronal enhancers fine-tune adaptive circuit plasticity. Neuron 2024;112:3043–3057.

[23] Han Q, Wang H, Lu X, Li Y, Guo Y, Zhao X, Feng Y, Hu L. Preoperative resting-state electrophysiological signals predict acute but not chronic postoperative pain. European Journal of Pain 2025;29:e4757.

[24] Hennessy SL, Sachs ME, Ilari B, Habibi A. Effects of Music Training on Inhibitory Control and Associated Neural Networks in School-Aged Children: A Longitudinal Study. Front Neurosci 2019;13. doi:10.3389/fnins.2019.01080.

[25] Herholz SC, Zatorre RJ. Musical training as a framework for brain plasticity: behavior, function, and structure. Neuron 2012;76:486–502.

[26] Hirano M, Kimoto Y, Furuya S. Specialized Somatosensory–Motor Integration Functions in Musicians. Cerebral Cortex 2020;30:1148–1158.

[27] Jensen MP, Gianas A, Sherlin LH, Howe JD. Pain Catastrophizing and EEG-α Asymmetry. The Clinical Journal of Pain 2015;31:852.

[28] Joret M-E, Germeys F, Gidron Y. Cognitive inhibitory control in children following early childhood music education. Musicae Scientiae 2017;21:303–315.

[29] Kim JA, Bosma RL, Hemington KS, Rogachov A, Osborne NR, Cheng JC, Oh J, Crawley AP, Dunkley BT, Davis KD. Neuropathic pain and pain interference are linked to alpha-band slowing and reduced beta-band magnetoencephalography activity within the dynamic pain connectome in patients with multiple sclerosis. PAIN 2019;160:187.

[30] Kleber B, Sitges C, Brattico E, Vuust P, Zamorano AM. Association Between Interoceptive Accuracy and Pain Perception: Insights From Trained Musicians and Athletes. European Journal of Pain 2025;29:e70012.

[31] Llinás RR, Ribary U, Jeanmonod D, Kronberg E, Mitra PP. Thalamocortical dysrhythmia: A neurological and neuropsychiatric syndrome characterized by magnetoencephalography. Proceedings of the National Academy of Sciences 1999;96:15222–15227.

[32] Luo C, Tu S, Peng Y, Gao S, Li J, Dong L, Li G, Lai Y, Li H, Yao D. Long-Term Effects of Musical Training and Functional Plasticity in Salience System. Neural Plasticity 2014;2014:1–13.

[33] Manning JR, Jacobs J, Fried I, Kahana MJ. Broadband Shifts in Local Field Potential Power Spectra Are Correlated with Single-Neuron Spiking in Humans. J Neurosci 2009;29:13613–13620.

[34] Mazaheri A, Seminowicz DA, Furman AJ. Peak alpha frequency as a candidate biomarker of pain sensitivity: the importance of distinguishing slow from slowing. NeuroImage 2022;262:119560.

[35] Millard SK, Furman AJ, Kerr A, Seminowicz DA, Gao F, Naidu BV, Mazaheri A. Predicting postoperative pain in lung cancer patients using preoperative peak alpha frequency. British Journal of Anaesthesia 2022;128:e346–e348.

[36] Oostenveld R, Fries P, Maris E, Schoffelen J-M. FieldTrip: Open Source Software for Advanced Analysis of MEG, EEG, and Invasive Electrophysiological Data. Computational Intelligence and Neuroscience 2011;2011:156869.

[37] Pascual-Leone A. The brain that plays music and is changed by it. Annals of the New York Academy of Sciences 2001;930:315–329.

[38] Pertermann M, Bluschke A, Roessner V, Beste C. The Modulation of Neural Noise Underlies the Effectiveness of Methylphenidate Treatment in Attention-Deficit/Hyperactivity Disorder. Biological Psychiatry: Cognitive Neuroscience and Neuroimaging 2019;4:743–750.

[39] Pertermann M, Mückschel M, Adelhöfer N, Ziemssen T, Beste C. On the interrelation of 1/f neural noise and norepinephrine system activity during motor response inhibition. Journal of Neurophysiology 2019;121:1633–1643.

[40] Ploner M, Sorg C, Gross J. Brain Rhythms of Pain. Trends in cognitive sciences 2017;21:100–110.

[41] Rosenkranz K, Williamon A, Rothwell JC. Motorcortical Excitability and Synaptic Plasticity Is Enhanced in Professional Musicians. J Neurosci 2007;27:5200–5206.

[42] Schaworonkow N, Voytek B. Longitudinal changes in aperiodic and periodic activity in electrophysiological recordings in the first seven months of life. Developmental Cognitive Neuroscience 2021;47:100895.

[43] Silva-Passadouro B, Delgado-Sanchez A, Henshaw J, Lopez-Diaz K, Trujillo-Barreto NJ, Jones AKP, Sivan M. Frontal alpha asymmetry: A potential biomarker of approach-withdrawal motivation towards pain. Front Pain Res 2022;3. doi:10.3389/fpain.2022.962722.

[44] Smith EE, Reznik SJ, Stewart JL, Allen JJB. Assessing and conceptualizing frontal EEG asymmetry: An updated primer on recording, processing, analyzing, and interpreting frontal alpha asymmetry. International Journal of Psychophysiology 2017;111:98–114.

[45] Stanhope J, Weinstein P. Should musicians play in pain? British Journal of Pain 2021;15:82–90.

[46] Valentini E, Halder S, McInnerney D, Cooke J, Gyimes IL, Romei V. Assessing the specificity of the relationship between brain alpha oscillations and tonic pain. NeuroImage 2022;255:119143.

[47] Vanneste S, De Ridder D. Chronic pain as a brain imbalance between pain input and pain suppression. Brain Communications 2021;3:fcab014.

[48] Voytek B, Kramer MA, Case J, Lepage KQ, Tempesta ZR, Knight RT, Gazzaley A. Age-Related Changes in 1/f Neural Electrophysiological Noise. J Neurosci 2015;35:13257–13265.

[49] de Vries M, Wilder-Smith OH, Jongsma ML, van den Broeke EN, Arns M, van Goor H, van Rijn CM. Altered resting state EEG in chronic pancreatitis patients: toward a marker for chronic pain. Journal of Pain Research 2013;6:815–824.

[50] Waschke L, Kloosterman NA, Obleser J, Garrett DD. Behavior needs neural variability. Neuron 2021;109:751–766.

[51] Zamorano AM, Cifre I, Montoya P, Riquelme I, Kleber B. Insula-based networks in professional musicians: Evidence for increased functional connectivity during resting state fMRI. Human Brain Mapping 2017;38:4834–4849.

[52] Zamorano AM, De Martino E, Insausti-Delgado A, Vuust P, Flor H, Graven-Nielsen T. Impact of Chronic Pain on Use-Dependent Plasticity: Corticomotor Excitability and Motor Representation in Musicians With and Without Pain. Brain Topogr 2024;37:874–880.

[53] Zamorano AM, Kleber B, Arguissain F, Boudreau S, Vuust P, Flor H, Graven-Nielsen T. Extensive Sensorimotor Training Predetermines Central Pain Changes During the Development of Prolonged Muscle Pain. The Journal of Pain 2023;24:1039–1055.

[54] Zamorano AM, Kleber B, De Martino E, Insausti-Delgado A, Vuust P, Flor H, Graven-Nielsen T. Prior use-dependent plasticity triggers different individual corticomotor responses during persistent musculoskeletal pain. PAIN 2025;166:e856.

[55] Zamorano AM, Kleber B, Martino ED, Insausti-Delgado A, Vuust P, Flor H, Graven-Nielsen T. Prior use-dependent plasticity triggers different individual corticomotor responses during persistent musculoskeletal pain. 2025:2025.01.15.633250. doi:10.1101/2025.01.15.633250.

[56] Zamorano AM, Montoya P, Cifre I, Vuust P, Riquelme I, Kleber B. Experience-dependent neuroplasticity in trained musicians modulates the effects of chronic pain on insula-based networks – A resting-state fMRI study. NeuroImage 2019;202:116103.

[57] Zamorano AM, Riquelme I, Kleber B, Altenmüller E, Hatem SM, Montoya P. Pain sensitivity and tactile spatial acuity are altered in healthy musicians as in chronic pain patients. Frontiers in Human Neuroscience 2015;8:1016.

[58] Zhang C, Stock A-K, Mückschel M, Hommel B, Beste C. Aperiodic neural activity reflects metacontrol. Cereb Cortex 2023;33:7941–7951.

[59] Zhang X, Bachmann P, Schilling TM, Naumann E, Schächinger H, Larra MF. Emotional stress regulation: The role of relative frontal alpha asymmetry in shaping the stress response. Biological Psychology 2018;138:231–239.

